# Multivariate pattern analysis for MEG: a comprehensive comparison of dissimilarity measures

**DOI:** 10.1101/172619

**Authors:** Matthias Guggenmos, Philipp Sterzer, Radoslaw Martin Cichy

## Abstract

Multivariate pattern analysis (MVPA) methods such as decoding and representational similarity analysis (RSA) are growing rapidly in popularity for the analysis of magnetoencephalography (MEG) data. However, little is known about the relative performance and characteristics of the specific dissimilarity measures used to describe differences between evoked activation patterns. Here we used a multisession MEG dataset to qualitatively characterize a range of dissimilarity measures and to quantitatively compare them with respect to classification accuracy (for decoding) and between-session reliability of representational dissimilarity matrices (for RSA). We tested dissimilarity measures from a range of classifiers (Linear Discriminant Analysis – LDA, Support Vector Machine – SVM, Weighted Robust Distance – WeiRD, Gaussian Naïve Bayes – GNB) and distances (Euclidean distance, Pearson correlation). In addition, we evaluated three key processing choices: 1) preprocessing (noise normalisation, removal of the pattern mean), 2) weighting classification accuracies by decision values, and 3) computing distances in three different partitioning schemes (non-cross-validated, cross-validated, within-class-corrected). Four main conclusions emerged from our results. First, multivariate noise normalization dramatically improved classification accuracies and the reliability of dissimilarity measures. Second, LDA, SVM and WeiRD yielded high peak classification accuracies and nearly identical time courses. Third, while using classification accuracies for RSA was markedly less reliable than continuous distances, this disadvantage was ameliorated by decision-value-weighting of classification accuracies. Fourth, the cross-validated Euclidean distance provided unbiased distance estimates and highly replicable representational dissimilarity matrices. Overall, we strongly advice the use of multivariate noise normalisation as a general preprocessing step, recommend LDA, SVM and WeiRD as classifiers for decoding and highlight the cross-validated Euclidean distance as a reliable and unbiased default choice for RSA.

**Highlights:**

1. We compared dissimilarity measures and preprocessing choices for MEG MVPA
2. Multivariate noise normalisation is a key preprocessing step
3. LDA, SVM and WeiRD are recommended classifiers for decoding
4. Use of decision values improves reliability of classification accuracies
5. The cross-validated Euclidean distance is a reliable and unbiased choice for RSA

## Introduction

The investigation of the rapid neural dynamics underlying cognitive functions requires a combination of high-temporal resolution neural measurements with analytical methods that systematically and efficiently probe the information encoded in measured brain activity. A promising approach is the application of multivariate pattern analysis methods (MVPA) to magnetoencephalography (MEG), combining the sensitivity of pattern-based methods with the high temporal resolution of MEG. Two prominent MVPA methods are multivariate decoding (Cox and Savoy, 2003; Haxby et al., 2001; Haynes and Rees, 2005; Kamitani and Tong, 2005), which quantifies the discriminability of condition-specific activation patterns, and representational similarity analysis (RSA) (Kriegeskorte et al., 2008). RSA characterizes the similarity of measured responses to experimental conditions in representational dissimilarity matrices (RDMs). As RDMs can in principle be computed for any measurement modality, RSA on MEG opens the way to quantitatively relate rapidly emerging brain dynamics to other data, such as fMRI (Cichy et al., 2016b, 2013) in order to localize responses; computational models (Cichy et al., 2017a, 2016a; Kietzmann et al., 2017; Pantazis et al., 2017; Seeliger et al., 2017; Su et al., 2012; Wardle et al., 2016) in order to understand the underlying algorithms and representational format; to behaviour (Cichy et al., 2017b; Furl et al., 2017; Mur et al., 2013); and across species (Cichy et al., 2014).

At the core of MVPA is the dissimilarity measure used to quantify the discriminability (decoding) or the dissimilarity structure (RSA) of evoked activation patterns, fundamentally affecting both the accuracy and the interpretability of results. Yet little is known about the performance and characteristics of different dissimilarity measures for MEG MVPA. Inspired by previous work comparing different dissimilarity measures for fMRI (Walther et al., 2016), we conducted a comprehensive and systematic investigation of dissimilarity measures for MEG to close this gap.

To this end, we compared a set of dissimilarity metrics comprising classifiers (Linear Discriminant Analysis - LDA, Support Vector Machine – LDA, Weighted Robust Distance – WeiRD, Gaussian Naïve Bayes – GNB) and distance measures (Euclidean distance, Pearson correlation). This comparison was done qualitatively, by characterizing “raw” dissimilarity time courses, and quantitatively, by comparing classification accuracies (decoding) and session-to-session reliabilities of RDMs (RSA). We further evaluated the effects of three main processing choices that affect dissimilarity estimation: 1) preprocessing (noise normalisation, removal of the pattern mean), 2) the use of classification decision values to preserve gradual information in classification-based MVPA, and 3) data partitioning (non-cross-validated; cross-validated; within-class-corrected, i.e. subtracting within- from between-condition distances).

Our results give rise to four straightforward recommendations for MVPA in MEG research. First, multivariate noise normalisation is strongly recommended as a general preprocessing step. Second, for decoding we recommend LDA, SVM and WeiRD, which achieved high accuracies. Third, we show that a previously reported impairment of pattern reliability for classification accuracy (Walther et al., 2016) can be mitigated by weighting correct and incorrect predictions with classifier decision values. Fourth and finally, concerning distance-based dissimilarity measures for RSA, we recommend the cross-validated Euclidean distance as a robust, gradual, reliable and largely unbiased default choice.

## Materials and Methods

### Dataset

The present study is based on a previously published MEG dataset (Cichy et al., 2014). This dataset was chosen for three reasons. First, the dataset is well powered in terms of the number of participants (N=16), the number of conditions (stimulus categories; N_c_=92) and the number of trials per condition (20–30 per session), enabling us to statistically evaluate even subtle effects of interest. Second, the dataset has two experimental sessions per participants, enabling us to compute inter-session reliabilities of our measures. Although it is possible to split a single experimental session into subparts to compute reliability, we reasoned that two independent sessions more realistically probe the robustness of a measure with respect to measurement quality (e.g., noise level of individual channels) or daily conditions of participants (e.g. wakefulness or motivation). Third, the employed stimulus set has been used in a number of previous studies (Cichy and Pantazis, 2016; Cichy et al., 2016b, 2014; Khaligh-Razavi and Kriegeskorte, 2014; Kiani et al., 2007; Kriegeskorte et al., 2008; Mur et al., 2013; Walther et al., 2016), facilitating the comparison of our results with previous literature.

We briefly summarize the most relevant aspects of experimental design and acquisition underlying the present dataset (for a detailed description, see Cichy et al., 2014). Participants viewed coloured images of 92 different objects on a grey background presented at the center of a screen (2.9° visual angle, 500 ms duration), overlaid with a dark grey fixation cross. For each of two MEG sessions, participants completed 10 to 15 runs of 420 s duration each. Each image was presented twice in each MEG run in random order, with a trial onset asynchrony of 1.5 or 2 s. To control vigilance and eye blink behaviour, participants were instructed to press a button and blink their eyes in response to a paper clip that was shown randomly every 3 to 5 trials (average 4). Paper clip trials were excluded from further analysis.

During the experiment, continuous MEG signals from 306 channels (204 planar gradiometers, 102 magnetometers, Elekta Neuromag TRIUX, Elekta, Stockholm) were acquired at a sampling rate of 1000 Hz. Recorded MEG signals were filtered between 0.03 and 330 Hz and preprocessed using spatiotemporal filters (maxfilter software, Elekta, Stockholm). Raw MEG trials were extracted with 100 ms baseline and a 1000 ms post-stimulus period (i.e., 1101 ms length), yielding 306-dimensional pattern vectors for each time point of a trial. In addition, raw trials were downsampled by averaging across consecutive 10 ms bins to decrease the computational costs and to increase the signal-to-noise ratio.

### Additional preprocessing of raw trials

Prior to MVPA, the MEG data may undergo additional preprocessing. Here, we assessed two popular preprocessing choices: 1) noise normalisation to improve the quality of the data, and 2) removal of the mean pattern to eliminate condition-nonspecific response components.

#### Multivariate noise normalisation

Commonly, MEG sensors differ in noise levels. To better exploit the information contained in multisensor MEG data, sensors with high noise levels (i.e., unreliable sensors) should be downweighted and sensors with low noise levels (i.e., reliable sensors) should be emphasized. This can be achieved by *univariate noise normalisation* (UNN), where each channel individually is normalised by an estimate of its error variance. Alternatively, it may be useful to emphasize or deemphasize specific spatial frequencies of MEG patterns. This can be achieved by means of *multivariate noise normalisation* (MNN), where in addition the error covariance between different sensors is considered. In both procedures, the MEG patterns **x** are normalised by means of a (co)variance matrix *Σ*:

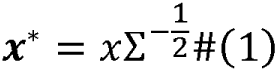

For MNN, off-diagonal elements of *Σ* correspond to the respective covariances, for UNN they are set to 0.

The covariance matrix *Σ* can be obtained in several different ways characterized by data selection (i.e., baseline phase or entire epoch) and by the level of temporal specificity (i.e., whether the covariance matrix is computed separately for each condition or each time point). To determine the best method for our dataset, in a first step we compared the performance of different covariance estimation methods. In brief, we computed *Σ* either on baseline data (*baseline method*), on the full epoch (*epoch method*) or separately for each time point (*time point method*). Moreover, since rank deficiency is often a problem for matrix inversion, we additionally tested a shrinkage transformation (Ledoit and Wolf, 2004) for the covariance matrix. The supplementary section “Comparison of noise normalisation methods” provides a detailed motivation and description of the methods.

Our main findings (summarized in Figure S1) were that 1) shrinkage improved the performance of all normalisation methods, 2) MNN was superior to UNN for the epoch and time point method and on par for the baseline method, and 3) the two overall best normalisation methods used MNN based on the epoch or the time point method. Given the slightly higher computational costs of the time point method, for the present work we chose MNN based on the epoch method.

#### Removing the mean pattern (cocktail-blank removal)

One likely complication when evaluating the dissimilarity of activation patterns is that different experimental conditions often share a common response component, irrespective of how different the conditions are. This has two main reasons. First, besides ample differences, experimental conditions may also have common experimental aspects, such as the fact that *any* stimulus was presented, leading to the activation of identical or overlapping neuronal populations. Second, even responses from non-overlapping, but spatially clustered, neuronal populations can produce similar activation patterns at the coarse spatial scale of neuroimaging measurements.

To account for condition-nonspecific response components, previous studies have subtracted the mean pattern from all conditions (“cocktail-blank removal”) (Op de Beeck, 2010; Pietrini et al., 2004; Williams et al., 2008, 2007). However, mean pattern removal is problematic in studies that delineate representational structure, such as RSA, as it introduces dependencies between conditions that may complicate the interpretability of RDMs (Diedrichsen et al., 2011; Garrido et al., 2013; Walther et al., 2016). Nevertheless, cocktail-blank removal is useful as a tool to determine the influence of condition-non-specific response patterns on the reliability of dissimilarity measures in case of RSA. In the present work, we used mean pattern removal only for this aim, assessing the effect of condition-nonspecific response patterns on the reliability of the Pearson distance in RSA (note that Euclidean distances are unaffected by condition-nonspecific patterns).

### Pseudo-trials

To increase the signal-to-noise ratio, for each of the N_c_ (=92) conditions we created 5 *pseudo*-*trials* from a random permutation of preprocessed raw trials (Figure 1A). This procedure was repeated for 20 permutations, such that in each permutation, raw trials of each condition were pseudo-randomly assigned to 5 pseudo-trials and then averaged.

**Figure 1.**
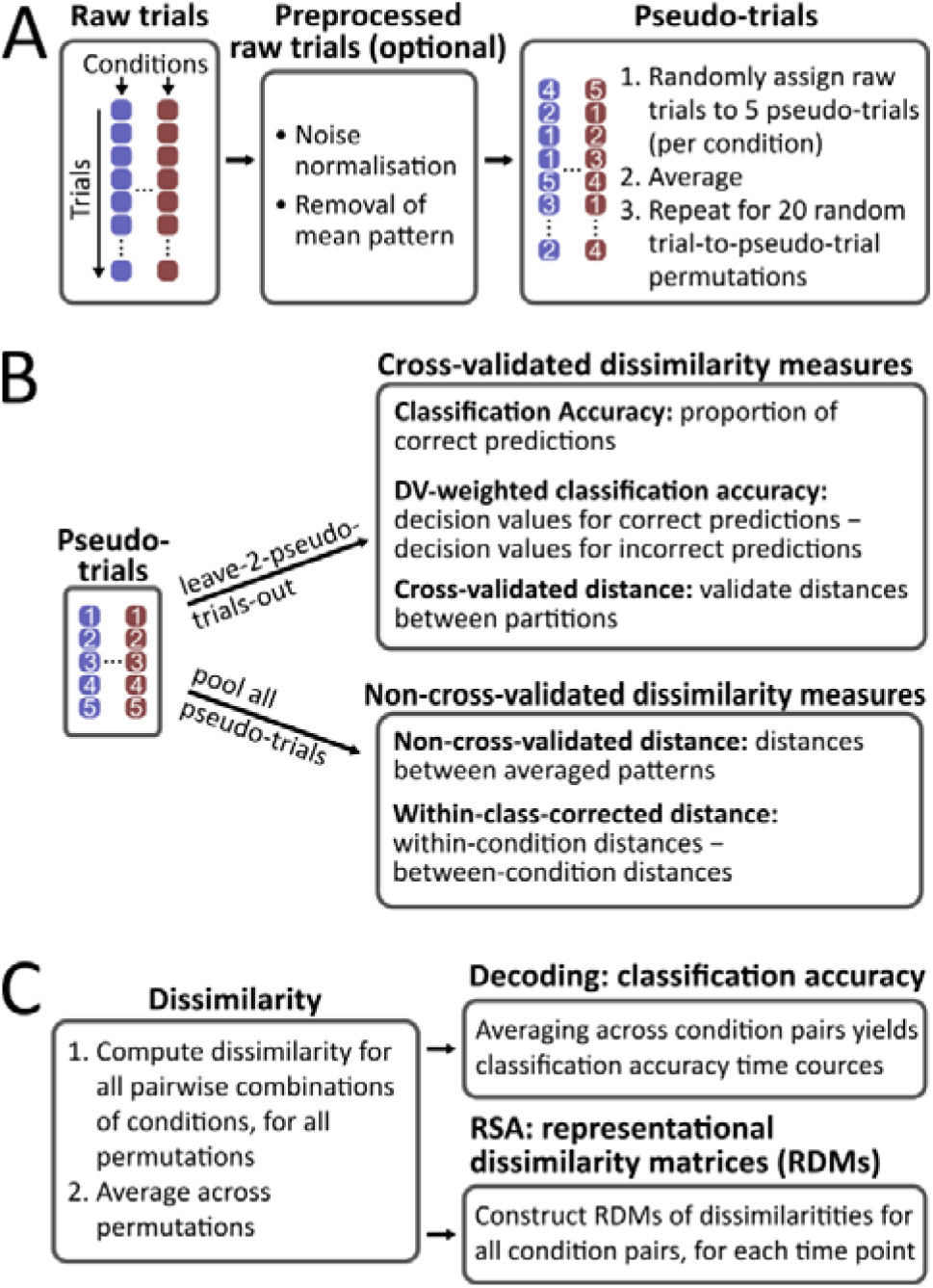
Analysis schemes and types of dissimilarity measures. **(A)** Preprocessing and pseudo-trials. After preprocessing, raw trials of each condition were assigned to 5 pseudo-trials through random permutation and averaged. **(B)** Overview of dissimilarity types for the computation of dissimilarity on pseudo-trials. Cross-validated measures are classification accuracy, decision-value(DV)-weighted classification accuracy and cross-validated distances; non-cross-validated measures comprise non-cross-validated and within-class-corrected distances. **(C)** Decoding and RSA. Condition-pair-specific dissimilarities are computed for 20 permutations (random assignments of trials to pseudo-trials) and then averaged across permutations. For decoding, classification accuracies were averaged across condition pairs. For RSA, N_c_ × N_c_ representational dissimilarity matrices (RDMs) were constructed for each time point.

### Data partitioning and cross-validation

Dissimilarity measurements between the pseudo trials of different condition was different depending on whether the employed dissimilarity measure was cross-validated or not (Figure 1B). For cross-validated measures (i.e., classification, cross-validated distances), the distance of two compared conditions was computed in a leave-two-(pseudo-trials)-out (LTO) cross-validation procedure. In each permutation, all but one randomly chosen pseudo-trial of each of any two compared conditions were used for training (hereafter referred to as data partition *A*) and the two left-out pseudo-trials were used for validation (data partition *B*). For non-cross-validated dissimilarity measures (i.e., non-cross-validated distances, within-class-corrected distances), all pseudo-trials of two compared conditions were pooled and the dissimilarity measure was computed once. More specifically, in the case of non-cross-validated distances, we averaged pseudo-trials and computed the distance based on this average. By contrast, because within-class-corrected distances required the distance computation to be carried out *within* a condition (i.e. between the individual pseudo-trials in a condition), pseudo-trials were in this case not averaged prior to distance computation.

### Types of dissimilarity measures

In this section, we introduce the dissimilarity measures compared in this work, which we divide into four different groups: 1) classification, 2) non-cross-validated, 3) cross-validated and 4) within-class-corrected distances.

In the following, vectors ***x*** and ***y*** represent the measured MEG patterns associated with two experimental conditions. Each pattern vector has length N_s_ corresponding to the number of sensors of the MEG device and is computed as the average vector for a considered condition and partition of the data. The goal is to compute a distance *d*(***x***, ***y***) between each pairwise combination of overall N_c_ conditions, ultimately resulting in a N_c_ × N_c_ representational dissimilarity matrix (RDM).

#### Classification accuracy

In classification, an algorithm is trained to discriminate between a set of conditions on the basis of labelled training samples. Here, we used binary classification, where a classifier is applied to each possible pairwise combination of the N_c_ conditions. Note that classification accuracies as reported here were always cross-validated, as in each fold classifiers were trained on one partition of the dataset (partition *A*) and tested on the left-out partition (partition *B*).

We assessed three common classifiers: Linear Discriminant Analysis classifier (LDA), linear Support Vector Machine (SVM) and Gaussian Naïve Bayes (GNB). We used the implementations provided by Scikit-learn (Abraham et al., 2014) with default parameters (in particular a cost parameter of 1 for SVM and the LIBSVM backend; Chang and Lin, 2011). Exploratory analyses, in which parameters were optimized in a nested cross-validation procedure, did not yield benefits for any of the classification results and were thus discarded. Note that for LDA, multivariate noise normalisation is originally an integral part of the algorithm itself. Thus, henceforth *LDA without multivariate noise normalisation* refers to the case where the covariance matrix in the LDA algorithm was set to the identity matrix (equivalent to a Euclidean distance-to-centroid classifier). In addition, we included the recently proposed Weighted Robust Distance (WeiRD; Guggenmos et al., 2016; https://github.com/m-guggenmos/weird) as a fourth classifier. WeiRD is a distance-to-centroid classifier, where Manhattan distances are computed in a statistically weighted feature space. A more detailed description is provided in the supplementary section “Weighted robust distance”.

The dissimilarity measure based on classification was *classification accuracy* and was defined as follows:

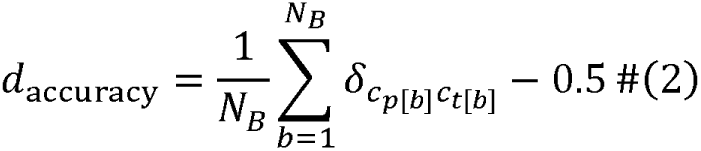

where *c_p_* is the condition predicted by the classifier, *c_t_* the true condition, *δ* the Kronecker delta, *N_B_* the number of samples in data partition *B*, and 0.5 the chance level. This definition of decoding accuracy has methodological advantages (expected value of zero in the absence of discriminative information; unitless) and is therefore used for all computations. However, note that for illustration purposes in figures we show classification accuracy as percent correct classification.

#### Decision-value-weighted classification accuracy

A possible drawback of classification accuracy as a distance metric in RSA is the loss of information due to the discretization into correct and incorrect predictions (Walther et al., 2016). Yet, all four classifiers construct some form of an internal continuous decision value (DV), which can be used to ameliorate the drawbacks of discretization. For the present set of classifiers, the decision value *DV* was either the absolute distance to a decision boundary (LDA, SVM, WeiRD) or the probability for the predicted class minus the chance level 0.5 (GNB).

The dissimilarity measure based on DV-weighted classification accuracies was defined as follows:

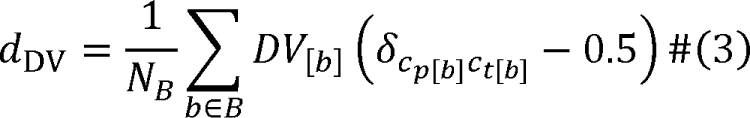

#### Non-cross-validated distances

Non-cross-validated distances do not require partitioning of the data and can be applied to patterns ***x*** and ***y*** averaged across all samples in a dataset. This has both advantages and disadvantages as compared to cross-validated distances. The advantages are that 1) the distance measure is often more robust because distances are computed on a larger set of data, and 2) the procedure is computationally more efficient, as the distance measure is computed only once for a comparison of two conditions. The main disadvantage is that non-cross-validated distances, in addition to estimating the true underlying distance, also capture the dissimilarity due to any source of noise. As a consequence, the measured distance can be biased by noise (see also, Walther et al., 2016).

Here we applied non-cross-validated distances after averaging the pattern vectors across all pseudo-trials of a condition. We evaluated two distance measures, the (squared) Euclidean and the Pearson distance, defined as follows:

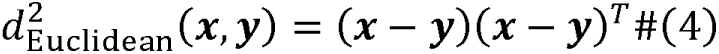

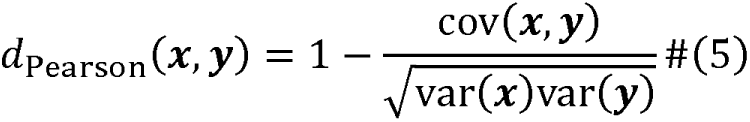

#### Cross-validated distances

To obtain unbiased measures of the true dissimilarity, as argued above it is necessary to compute distances in a cross-validated procedure. For this, the pattern vectors of one data partition are projected on those of an independent (validation) partition. As the noise components of the pattern vectors can be assumed to be mostly orthogonal across partitions, they become eliminated by cross-validation (mathematically, note that the projection product of orthogonal vectors is zero).

This property of cross-validation has two key advantages. First, it can improve the reliability of distance estimates when noise levels differ between measurements. Second, unbiased distances enable testing of ratio-based hypotheses, such as whether one distance is twice as big as another distance, or whether a distance is different from zero (Walther et al., 2016). Such a test would not be sensible if distances are affected by noise as in the case of non-cross-validated distances or classification.

Equations (6) and (7) define the *cross*-*validated (c.v.)* variants of the (squared) Euclidean and the Pearson distance:

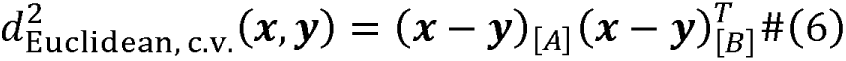

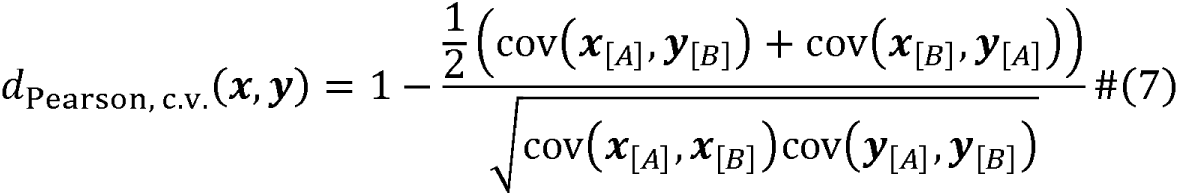

where A and B denote the two partitions of the data within each cross-validation fold (A: base partition, comprising all but 2 left-out pseudo-trials; B: validation partition, the 2 left-out pseudo-trials).

A practical issue for cross-validated Pearson distances is the fact that the variances in the denominator can be very small or even negative, leading to exceedingly negative, exceedingly positive or imaginary distances. To accommodate this issue, we regularized the cross-validated Pearson distance in three ways. First, we enforced a positive lower bound ∈ for the variance. As a sensible lower bound depends on the scaling of the data, the parameter ∈ was set to 10% of the non-cross-validated variance of data partition A (the value of 10% was determined from an independent simulation). Second, we enforced a lower bound for the denominator, for which the minimum value was set to 25% of the non-cross-validated denominator (likewise determined via simulation). And third, we bounded the resulting Pearson distance between 0 and 2 (i.e., corresponding to the bounds of the non-cross-validated Pearson distance).

#### Within-class-corrected distances

As outlined above, condition-nonspecific response patterns are a ubiquitous phenomenon in neuroimaging measurements. Critically, such condition-nonspecific patterns can impair the accuracy of RSA when they differentially affect specific conditions in two compared modalities (e.g., MEG and behaviour). Here we propose, as a remedy for condition-nonspecific patterns, a procedure we refer to as *within*-*class correction*, where within-condition dissimilarities are subtracted from between-condition dissimilarities. Within-class correction has been previously used to estimate the discriminatory power of activation patterns (Golarai et al., 2007; Haxby et al., 2001; Weiner et al., 2010). In the context of RSA, it may provide a solution to remove condition-nonspecific patterns in the computation of distance-based dissimilarities.

In addition to removing condition-nonspecific responses, within-class correction can yield unbiased distance estimates under certain circumstances. However, for the two distance measures under investigation this applies only to the Euclidean distance. By contrast, the within-class-corrected Pearson distance is not unbiased in the presence of noise, as both within- and between-condition distances approach 1 with increasing noise and the within-class-corrected distance thus 0. For the Euclidean distance, within-class correction yields an unbiased measure only if the within-condition noise is at the same level as between-condition noise. The fact that within-class correction is thus not generally unbiased is a disadvantage compared to cross-validation.

Mathematically, for the *within*-*class*-*corrected distances (w*.*c*.*c*.) we subtract the average within-condition distance from the between-condition distance:

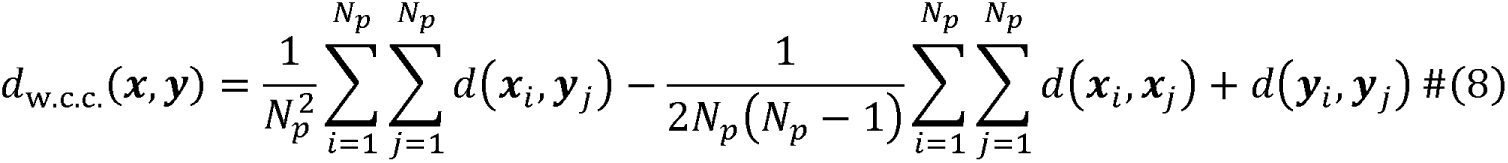

where *d* is the given distance measure and *N_p_* = 5 is the number of pseudo-trials of a condition. While the first double sum represents the average distance of all pairwise pseudo-trial combinations *between* the two compared conditions (between-condition distance), the second and subtracted double sum represents the average distance of all pairwise pseudo-trial combinations *within* each of the two conditions (within-condition distance).

### Decoding

Decoding uses classification to quantify the discriminability of activation patterns pertaining to a set of experimental conditions (Cox and Savoy, 2003; Haxby et al., 2001; Haynes and Rees, 2005; Kamitani and Tong, 2005). Here, we computed classification accuracies for all pairwise combinations of conditions and all permutations and then averaged across permutations and conditions, separately for each time point. As a result we obtained a single average classification accuracy time course for each participant.

### Representational similarity analysis

The goal of representational similarity analysis (RSA; Kriegeskorte et al., 2008) applied to neuroimaging data is to characterize the (dis)similarity of activation patterns for a set of experimental conditions. The dissimilarities between all pairwise combinations of conditions are organized in representational dissimilarity matrices (RDMs). In the present work, to construct RDMs we computed dissimilarities for all pairwise combinations of conditions and all permutations and then averaged across permutations, separately for each time point. As a result we obtained N_c_ × N_c_ RDMs for each time point and participant.

### Reliability measures

While classification accuracies can be directly compared to find the best classifier for decoding, such a direct comparison is not possible across the diverse full set of dissimilarity measures (classification accuracies, decision values or distances) that can be used to construct RDMs in RSA. For this reason, we used the session-to-session reliability of RDMs as a performance measure that generalizes across types of dissimilarity measures. The rationale for using reliability was that more robust and faithful dissimilarity measures show more replicable RDMs across sessions.

Here, we computed reliabilities between sessions, exploiting the fact that the dataset included two equivalent experimental sessions for each participant. In the following, we consider two types of RDM reliability.

First, using the Pearson correlation coefficient, we computed the strength of a linear relationship between two RDMs, regardless of mean and scaling. Mean and scale invariance can be desired properties, e.g. when the mean of two RDMs differs due to varying noise levels or when the scaling differs due to different transfer functions of the involved measurement devices. The *pattern reliability* was defined as follows:

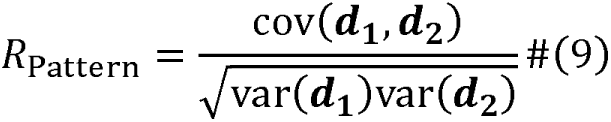

where ***d***_1_ and ***d***_2_ represent vectorised RDMs (i.e., the flattened lower triangular part of an RDM) pertaining to the two sessions. *R_Pattern_* can take values between –1 and 1.

On the other hand, it may often be deemed important that two RDMs are as close as possible in terms of their Euclidean distance, thus respecting both mean value and scaling. For instance, one may have reasons to believe that mean differences between RDMs reflect truthful differences that are not trivially explained by different noise levels (e.g., when noise levels were controlled or cross-validation was performed). We therefore additionally computed the normalised sum of squared differences (SSQ) between two RDMs, referred to as the *SSQ reliability* (Walther et al., 2016):

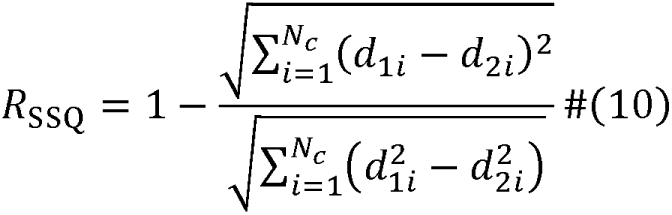

where *R_SSQ_* take values between 0 and 1.

Note that for the non-cross- and cross-validated Pearson distance, we computed the SSQ reliability on 1–d_1i_ and l–d_2i_ in order to enforce an expected dissimilarity of zero for random activation patterns, and thus an expected SSQ reliability of zero. This ensured comparability of the Pearson distance with other dissimilarity measures in terms of SSQ reliability.

Non-cross-validated Euclidean distances likewise do not lead to a distance of zero for random activation patterns. However, since this is a fundamental property of the noise dependency, rather than a technicality as whether to use d or 1–d, we omitted the non-cross-validated Euclidean distance when reporting SSQ reliability.

### Statistical testing

Statistical tests were applied to compare dissimilarity measures either in a time-resolved manner or averaged across time points within a time window of interest. In both cases, sign permutation tests were performed against the null hypothesis of no difference. We employed a full sign permutation scheme across 16 participants. The minimal possible *p*-value was thus 1/2^16^ = 1.526⋅10^−5^, which is denoted as *p* <2^−16^ in the text. For time-resolved statistical tests, sign permutation tests were used with Bonferroni-correction for the number of time points.

### Code

This article is accompanied by an online tutorial with code (Python, Matlab) and instructions to reproduce key results, available at https://github.com/m-guggenmos/megmvpa.

## Results

### 1 Classification accuracy: finding the best classifier for decoding

Overall, four different classifiers (LDA, SVM, WeiRD and GNB) with and without multivariate noise normalisation (MNN) were evaluated, where each classifier predicted stimulus category labels from MEG data in a time-resolved fashion. Time courses of classification accuracy were computed by comparison with true category labels (Figure 2).

**Figure 2.**
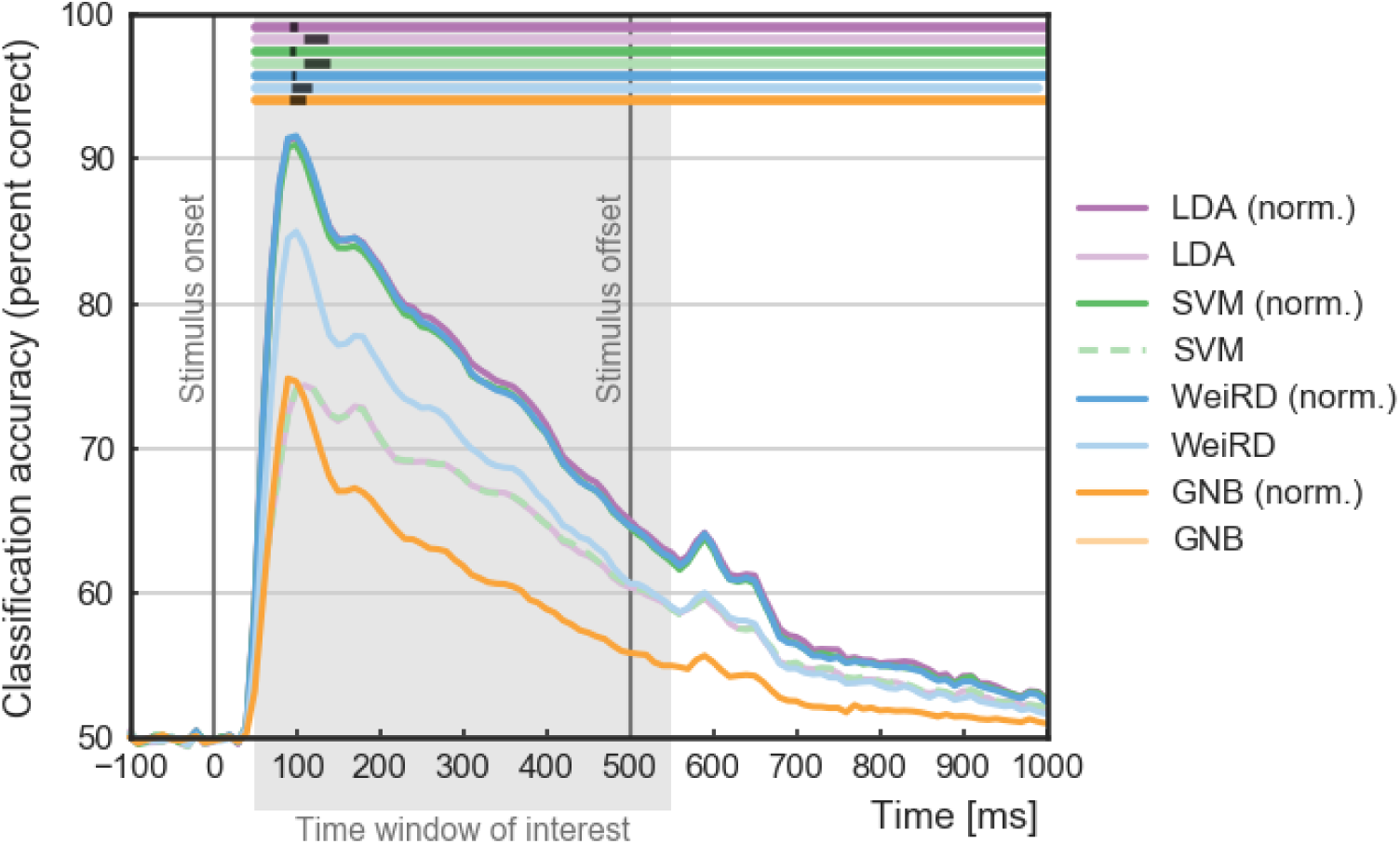
Decoding: time-resolved MEG classification accuracies. Color-coded curves (as in legend) report classification accuracy (percent correct) for the tested set of classifiers (LDA, SVM, WeiRD, GNB) with and without multivariate noise normalisation (norm.). Horizontal lines at the top of the figure mark statistically significant time points (sign permutation test, Bonferroni-corrected for the number of time points (100), corrected significance level *p*< 0.05). Semi-transparent black bars indicate the bootstrapped 95% confidence interval for peaks (10^6^ samples with replacement).

An analysis of classification accuracy time courses showed that the accuracy was above chance for all tested classifiers at each time point of the time window of interest (*p* < 0.05, sign permutation test, Bonferroni-corrected for 100 time points). The accuracy curves peaked around 100 ms (95 % confidence intervals illustrated as black bars in Figure 3 over curves) and decreased afterwards.

**Figure 3.**
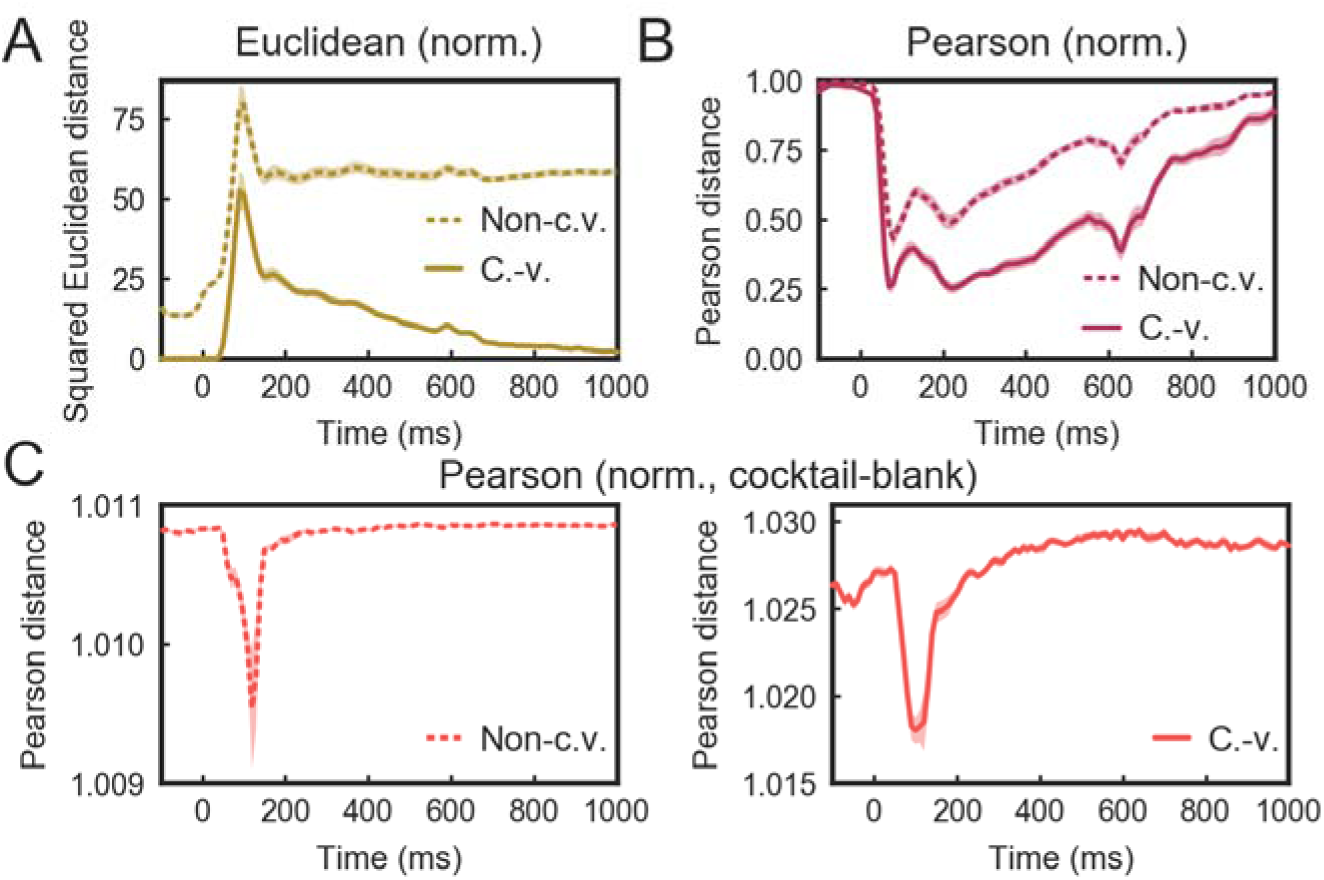
Raw distance measures across time with and without cross-validation. **(A)** Euclidean distance. **(B)** Pearson distance. **(C)** Pearson distance with cocktail-blank removal. Note that the Pearson distance exceeded a value of 1, which is indicative of a negative bias caused by cocktail-blank removal. For brevity and clarity, all distance measures are only shown with prior multivariate noise normalisation of the data. Shaded areas indicate bootstrapped standard errors across participants (10^5^ samples with replacement).

For a summary statistical comparison of classifiers, we averaged accuracies across time (from 50 to 550 ms, i.e. accounting for an ~50ms offset between stimulus presentation and cerebral processing). In a first step, we investigated the effect of MNN and found that applying MNN prior to classification boosted classification performance between 5 and 20% percent correct classification. This makes MNN an indispensable preprocessing step. Next, we compared classification accuracy across noise-normalised classifiers. Noise-normalised LDA, SVM and WeiRD performed comparably well with peak accuracies of over 90% correct classification, while GNB performed markedly worse (Figure S2). Thus, LDA, WeiRD and SVM with MNN were suitable choices for MEG pattern classification.

In conclusion, MNN is a highly recommended preprocessing step prior to classification. In terms of classifiers, our results indicate that LDA, SVM and WeiRD are all suitable and powerful classifiers and are thus recommended for multivariate decoding in MEG research.

### 2 Distance measures: characteristics and the effect of cross-validation and within-class correction

Before investigating reliabilities, it is helpful to identify special properties of different distance measures, as those impact the proper interpretation of session-to-session reliabilities. We thus begin our analysis with a characterization of raw dissimilarity time courses. Readers familiar with these properties or only interested in reliability may skip this section.

#### 2.1 Euclidean distance: noise dependency and a remedy through cross-validation

Without cross-validation, the non-cross-validated Euclidean distance (Figure 3A, dashed yellow curve) remained at a relatively high level from around 100 ms onwards, even though our classification analysis indicated a decrease of informative category-related signals during these later time points. How does cross-validation affect the Euclidean distance estimates across time? We found that cross-validation led to a gradual decrease of Euclidean distance estimates after 100 ms (Figure 3A, solid yellow curve), consistent with the decrease in classification accuracy. This result confirms the notion that cross-validation yields unbiased distance measures in the presence of noise (Walther et al., 2016) and thus addresses the noise inflation observed without cross-validation.

#### 2.2 Pearson distance: the effect of cross-validation and the critical issue of condition-nonspecific signal influences

Different from the Euclidean distance, the Pearson distance between entirely random patterns has an expected value of 1, regardless of the general noise level. Thus, already without cross-validation, the Pearson distance has a meaningful zero point (i.e., when subtracting a constant value of 1). Yet, the Pearson distance is by no means unaffected by noise: as correlation coefficients measuring non-zero correlations decrease with increasing noise, the measured Pearson distance will always show a smaller deviation from 1 compared to the true Pearson distance. This raises the question whether cross-validation, just like in the case of the Euclidean distance, could enable more faithful estimates of the Pearson metric.

The results for our data conformed to these expectations. During the baseline phase (0 to 100 ms) for which we can assume random MEG patterns, the Pearson distance was 1 both without and with cross-validation (Figure 3B). With the onset of the stimulus, the cross-validated Pearson distance (solid curve) became markedly lower than the non-cross-validated Pearson distance (dashed curve). This indicates that the non-cross-validated Pearson distance underestimated the true (positive) correlation between MEG patterns, a noise bias that was moderated by cross-validation.

Both without and with cross-validation, the time course of the Pearson distance showed a shape that may seem surprising at first: Pearson distances became *smaller* after stimulus onset, i.e. the patterns of different conditions became *more similar*. Indeed, the Pearson distance time courses showed an almost inverted shape compared to the cross-validated Euclidean distance or classification accuracies. The reason is that, although different conditions differed in stimulus content, they nevertheless shared a number of commonalities, not least the fact that *any stimulus* was presented or that there was always a *stimulus offset* (peak between 600 and 700ms). However, when activation patterns are rendered highly *similar* through a dominance of condition-nonspecific signals, the goal of RSA is at risk, i.e. mapping the *specific dissimilarity* structure between conditions.

To correct for condition-nonspecific signals, we subtracted the mean pattern from all conditions (cocktail-blank removal) for the Pearson distance (Figure 3C). We found that cocktail-blank removal was indeed successful in removing the bulk of condition-nonspecific signal contributions. However, confirming earlier work (Diedrichsen et al., 2011; Garrido et al., 2013; Walther et al., 2016), cocktail-blank removal led to negative correlations between activation patterns of conditions (i.e., Pearson distances greater than 1), even in the baseline phase. Because of these artefactual dependencies, in the present work we limited the application of cocktail-blank removal to assessing a potential inflationary effect of condition-nonspecific response patterns on RDM reliability.

#### 2.3 Within-class correction reveals the condition-specific component of non-cross-validated distances

Like cocktail-blank removal, the goal of *within*-*class correction* is to eliminate condition-nonspecific components from activation patterns (e.g., Haxby et al., 2001). This is achieved by subtracting within-condition distances from between-condition distances. The underlying premise is that condition-nonspecific signals not only affect distances between *different* conditions, but also between repeated measurements of the *same* condition. Within-class correction thus removes signal and noise components unrelated to the difference between conditions.

We found that, when using within-class correction, the raw dissimilarity time courses based on the Euclidean and Pearson distance metrics were more similar to the time courses of classification accuracy and thus had closer bearing to the discriminatory power of condition-specific activation patterns (Figure 4). For the Euclidean distance, within-class correction largely eliminated the substantial *condition*-*nonspecific noise component* that caused high Euclidean distances long after stimulus onset (Figure 4A). In contrast, for the Pearson distance, within-class correction removed the previously dominating *condition*-*nonspecific signal component* (Figure 4B). In particular, within-class correction revealed that the Pearson distance of condition-specific signal components indeed increased with stimulus presentation, just as in the case of the Euclidean distance. We thus conclude that within-class correction accounted for the effect of *condition*-*nonspecific response* components on dissimilarity, which otherwise affected the non-cross-validated Euclidean and Pearson distance as well as the cross-validated Pearson distance.

**Figure 4.**
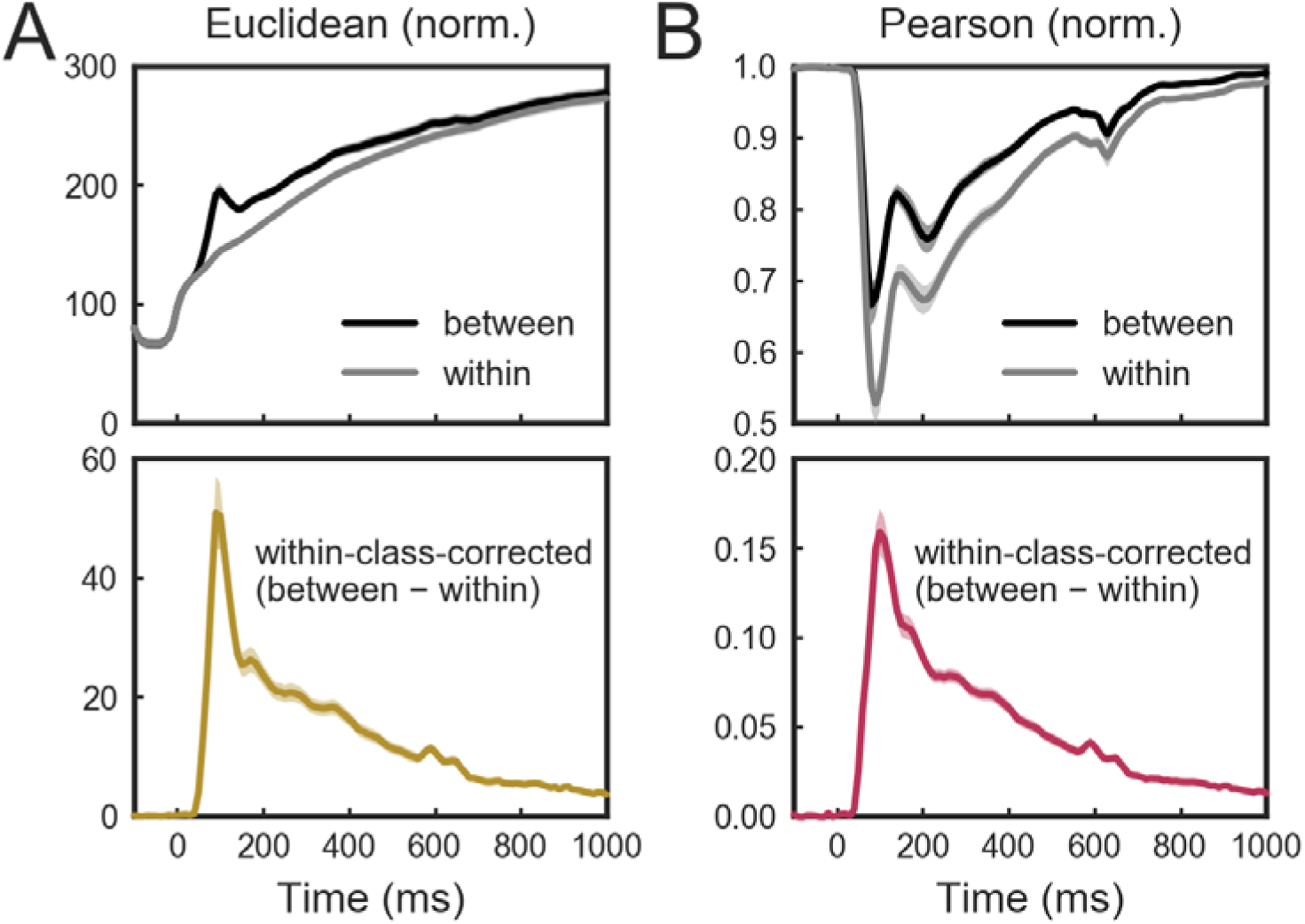
Within-class-correction of non-cross-validated distances. Top panels show within- and between-condition distances, bottom panels show difference curves (between minus within, referred to as within-class-corrected). Multivariately noise normalised **(A)** Euclidean distance and **(B)** Pearson distance. Shaded areas indicate bootstrapped standard errors across participants (10^5^ samples with replacement).

### 3 Reliability: finding the best dissimilarity measure for representational similarity analysis

We used the reliability of RDMs across measurements to compare the performance of all dissimilarity measures under investigation. Two different reliability measures were applied: *pattern reliability*, a correlation-based measure assessing pattern similarity irrespective of mean and scaling, and *SSQ reliability*, a Euclidean-distance-based measure assessing the similarity taking mean and scaling into account.

We report reliability time courses for all investigated dissimilarity measures: classification accuracies (Figure 5A), DV-weighted classification accuracies (Figure 5B), non-cross-validated distances (Figure 5C), cross-validated distances (Figure 5D) and within-class-corrected distances (Figure 5E). For summary purposes, we additionally computed the average and the maximum reliability over the time window of interest (Figure 6; Figure S3 for direct comparisons).

**Figure 5.**
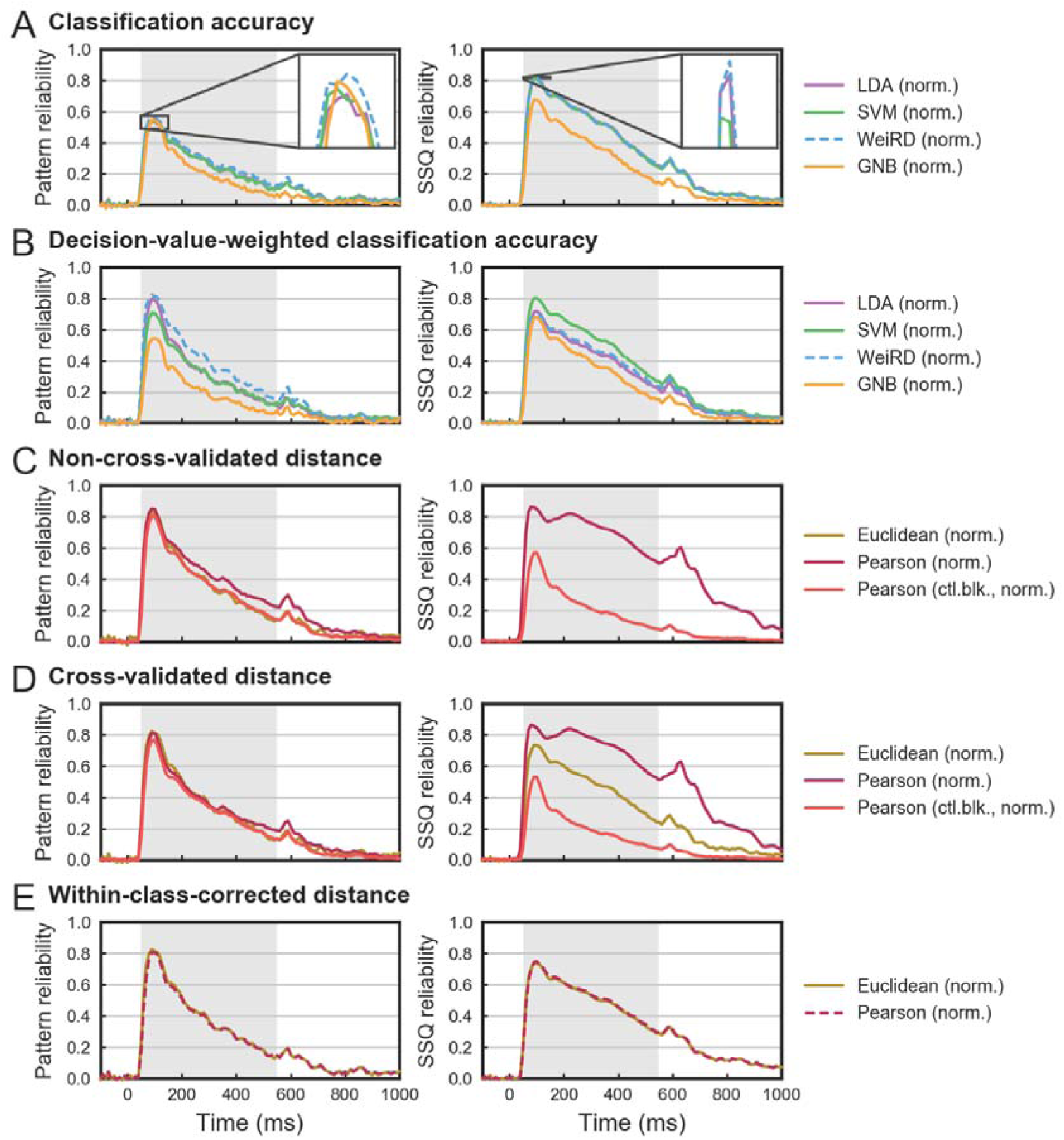
Time-resolved pattern and SSQ reliability of RDMs across sessions. For clarity, all measures are depicted only with multivariate noise normalisation. The shaded grey area depicts the time window of interest [50 ms; 550 ms]. **(A)** Classification accuracies. **(B)** Decision-value-weighted classification accuracies. **(C)** Non-cross-validated distances. Note that for the Euclidean distance, the SSQ reliability cannot be meaningfully computed and is therefore omitted. **(D)** Cross-validated distances. **(E)** Within-class-corrected distances. Abbreviations: norm. = multivariate noise normalisation; ctl.blk. = cocktail-blank removal

**Figure 6.**
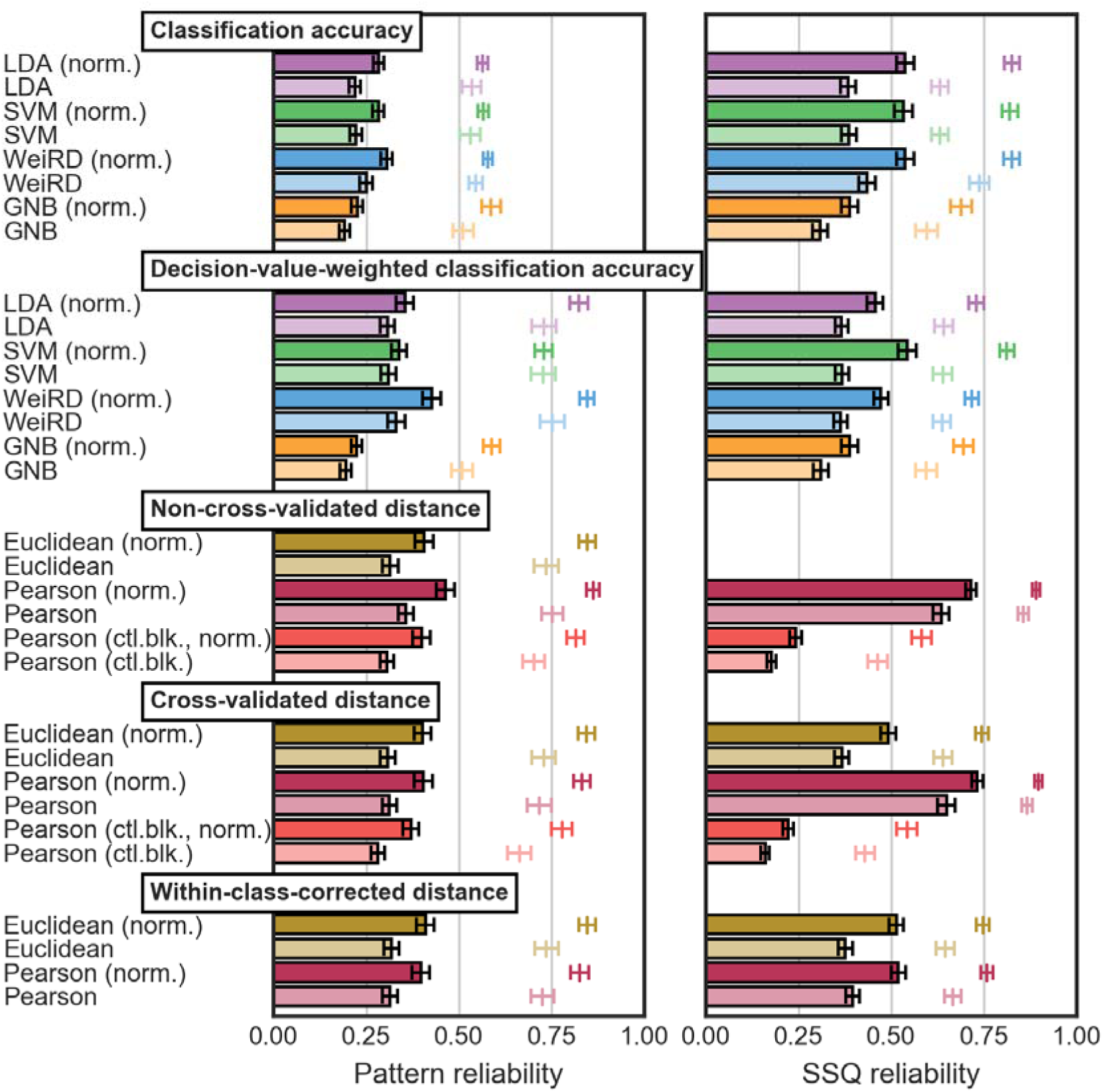
Average and maximum reliability of RDMs across sessions. Bars represent the average reliability, free-floating lines the maximum reliability within the time window [50 ms; 550 ms]. Error bars indicate bootstrapped standard errors across participants (10^6^ bootstrap samples). Abbreviations: norm. = multivariate noise normalisation; ctl.blk. = cocktail-blank removal

#### 3.1 Multivariate noise normalisation improves reliability

MNN substantially improved the reliability of all dissimilarity measures. Indeed, in most cases the choice of a dissimilarity measure itself was less critical than whether MNN was performed or not (Figure 6). Across dissimilarity measures, MNN led to an average gain of ΔR_Pattern_ = 0.07 and ΔR_SSQ_ = 0.11. In view of these generally beneficial effect of MNN on reliability, below we restricted all further analyses to cases where MNN was applied.

#### 3.2 The most accurate classifiers are also the most reliable classifiers

We showed earlier that LDA, SVM and WeiRD were comparable in accuracy, while GNB performed markedly worse. We found that this pattern of results was matched by our analysis of reliability. LDA, SVM and WeiRD showed only small reliability differences, while GNB fell off by a large margin. Between LDA, SVM and WeiRD, WeiRD slightly outperformed LDA (ΔR_Pattern_ = 0.023, *p* < 2^−16^, sign permutation test) and SVM (ΔR_Pattern_ = 0.023, *p* < 2^−16^) in terms of pattern reliability and SVM in terms of SSQ reliability (ΔR_SSQ_ = 0.004, *p* = 0.036); all other comparison were not significant.

We conclude that LDA, SVM and WeiRD were all suitable choices with respect to RDM reliability. In our dataset, WeiRD was slightly more reliable than LDA and SVM. More generally, the fact that the reliability analysis yielded much the same conclusion as a comparison of classification accuracies adds validation to the rationale of using reliability as a performance criterion for dissimilarity measures.

#### 3.3 The non-cross-validated Euclidean and Pearson distance are equally reliable when accounting for condition-nonspecific response components

Judging between the Euclidean and the Pearson distance, which is the more reliable distance measure when non-cross-validated, i.e. without cross-validation or within-class correction? We found that there are two answers to this question.

When considering raw pattern reliability^1^, the Pearson distance outperformed the Euclidean distance by a large margin (ΔR_Pattern_ = 0.057, *p* < 2^−16^). Indeed, in this view, the non-cross-validated Pearson distance was the most reliable measure across all tested measures.

However, given the strong contribution of condition-nonspecific response components to the Pearson distance, the question arises to what degree these shared components might inflate the reliability. With respect to the goals of RSA this is a critical question, because reliability gains based on response components that are shared between conditions are of no use. Evaluating the non-cross-validated Pearson distance after cocktail-blank removal revealed that the reliability of the non-cross-validated Pearson distance became indistinguishable from the non-cross-validated Euclidean distance (ΔR_Pattern_ = –0.007, *p* = 0.17). This suggests that the reliability advantage of the non-cross-validated Pearson distance was based on condition-nonspecific response components.

In sum, the non-cross-validated Euclidean and Pearson distance are equally reliable when condition-nonspecific response components are subtracted through cocktail-blank removal.

#### 3.4 Cross-validation can impair the reliability of distance measures

Two aspects of cross-validation may impact reliability in opposite ways. On the one hand, cross-validation provides faithful distance measures that are largely unbiased in the presence of noise. This could benefit reliability in cases where noise levels differ between measurements. On the other hand, the split of the data into cross-validation folds might negatively impact the accuracy of the estimated distance measure and thus reliability.

We found a marginal reduction in pattern reliability when cross-validating the Euclidean distance as compared to no cross-validation (ΔR_Pattern_ = –0.005, *p* = 0.068) and a significant loss in case of the Pearson distance (without cocktail-blank: ΔR_Pattern_ = –0.059, *p* < 2^−16^; with cocktail-blank: ΔR_Pattern_ = –0.029, *p* < 2^−16^). In terms of SSQ reliability, cross-validation had a negative effect on the Pearson distance without (ΔR_SSQ_ = –0.017, *p* = 0.0001), but not with cocktail-blank removal (ΔR_SSQ_ = 0.021, *p* < 2^−16^). These largely negative effects of cross-validation on reliability were confirmed by simulation (Figure S4), suggesting that they were generic and not specific to our datasets.

Taken together, while cross-validation has a unique advantage in terms of providing unbiased distance estimates, it may come at the cost of reduced reliability. In our empirical data, this cost was minor for the Euclidean distance and more pronounced for the Pearson distance.

#### 3.5 Within-class correction provides condition-specific estimates of the Pearson distance at high reliability

Within-class correction addresses the issue of condition-nonspecific signal and noise components that are shared between activation patterns of different conditions. Yet, how reliable are the RDMs generated through within-class-corrected distances compared to non-cross- and cross-validated distances?

For the Euclidean distance, we found equal reliability of within-corrected distances and non-cross-validated distances (ΔR_Pattern_ = 0.001, *p* = 0.33) and a slight advantage of within-class correction over cross-validation (ΔR_Pattern_ = -0.007, *p* = 0.0002; ΔR_SSQ_ = -0.022, *p* < 2^−16^). Thus, in terms of reliability only, within-class correction of the Euclidean distance is not associated with large changes in reliability.

For the Pearson distance, the effect of within-class correction strongly depended on whether non-cross- and cross-validated distances were computed with prior cocktail-blank removal. Without cocktail-blank removal, within-class-corrected distances were generally inferior to non-cross-validated (ΔR_Pattern_=0.066, *p* < 2^−16^; ΔR_SSQ_=0.196, *p* < 2^−16^) and cross-validated distances (ΔR_pattern_ = 0.007, *p*=n. s.; ΔRSSQ=0.213, *p* < 2 ^−16^).

By contrast, when the within-class-corrected Pearson distance was compared to the cocktail-blank-corrected non-cross- and cross-validated distances this pattern was largely reversed. Here, within-class-corrected distances showed strong reliability advantages compared to non-cross-validated (ΔR_SSQ_= –0.277, *p* < 2^−16^; but ΔR_pattern_ = 0.002, *p* = 0.33) and cross-validated (ΔR_pattern_ = –0.027, *p* = 0.0004; ΔR_SSQ_= –0.299, *p* < 2^−16^) distances. Thus, not only did within-class correction rectify the effect of condition-nonspecific response components on the Pearson distance in a more valid manner than cocktail-blank removal, it also did so at higher reliability.

Overall, our data showed that within-class correction had a small but positive influence on the reliability of Euclidean distances, and a strong effect on the reliability of the Pearson distances when those were corrected for condition-nonspecific response components. Of note, this pattern of results was confirmed by simulation (Figure S4). As a main conclusion, we recommend to use within-class correction for the Pearson distances when condition-nonspecific responses are deemed problematic.

#### 3.6 Decision-value weighting narrows the gap in pattern reliability between classification-based and distance-based RSA

Disadvantages of classification accuracies for RSA are that 1) gradual information contained in the decision values of classifier predictions is lost by using a binary measure for the *correctness* of predictions, and 2) classification accuracies naturally decrease with noise and are thus not unbiased. Walther et al. (2016) previously showed that these disadvantages can substantially impair pattern reliability of fMRI data. We observed results consistent with this prediction for MEG, i.e. classification accuracies exhibited generally lower pattern reliability than distances (Figure 6). Notably this pattern was reversed for SSQ reliability, where classification accuracies generally outperformed distance measures. Ironically, this is likely also due to the discretization of classification accuracies. Discretization constrains individual dissimilarity estimates to –0.5 and 0.5, effectively regularizing SSQ differences in preventing unrealistically large SSQ differences between activation patterns. Nevertheless, as the impairment of pattern reliability for classification accuracies is substantial, it should be avoided for RSA.

A potential remedy is to weigh the correctness of the predictions by decision values (DVs) (see Fig. S5 for raw time courses of DV-weighted classification accuracies). We found that for most classifiers, DV-weighting substantially increased the pattern reliability compared to raw classification accuracies (Figure 6B). Yet, DV-weighting impaired the SSQ reliability for most classifiers (exception: SVM), likely because decision values were much more volatile and prone to extreme values than classification accuracies, which are constrained between –0.5 and 0.5 (i.e., 0 and 100% in percent correct). Thus, while DV-weighting can lead to considerable improvements in pattern reliability, this may have to be traded off against an impairment in SSQ reliability.

Across classifiers, we found that the pattern reliability of DV-weighted classification accuracies was higher for WeiRD than LDA (ΔR_Pattern_ = 0.072, *p* < 2^−16^) and SVM (ΔR_Pattern_ = 0.088, p < 2^−16^), and higher for LDA than SVM (ΔR_Pattern_ = 0.016, *p* =0.028). In terms of SSQ reliability, SVM outperformed LDA (ΔR_ssq_= 0.087, *p* < 2^−16^) and WeiRD (ΔR_SSQ_ = 0.071, p < 2^−16^), and WeiRD outperformed LDA (ΔR_SSQ_ = 0.022, *p* = 0.012). Thus, when classification-based dissimilarity measures are desired for RSA, DV-weighted classification accuracies of WeiRD and SVM may be particularly attractive choices depending on whether one prioritizes pattern and SSQ reliability, respectively.

How does the reliability of DV-weighted classification accuracies compare with distance measures? We here report the comparison with the distance schemes for the Euclidean and Pearson metric that we recommended based on the results in the previous two sections: cross-validated Euclidean distances and within-class-corrected Pearson distances. Our results show that LDA and SVM, despite the improvements in pattern reliability through DV-weighting, could not fully reach the pattern reliability of these two distance measures. However, DV-weighting of WeiRD classification accuracies yielded higher pattern reliability than both the cross-validated Euclidean distance (ΔR_Pattern_= 0.025, *p* < 2^−16^) and the within-class-corrected Pearson distance (ΔR_Pattern_ = 0.029, *p* < 2^−16^). In terms of SSQ reliability, while DV-weighting caused a lower SSQ reliability for WeiRD and LDA compared to the reference distances, the SSQ reliability of SVM became superior to the cross-validated Euclidean distance (ΔR_Pattern_= 0.052 *p* < 2^−16^) and the within-class-corrected Pearson distance (ΔR_Pattern_= 0.024, *p*= 0.0001). Thus, DV-weighting narrows the gap in pattern reliability between classification- and distance-based dissimilarity measures, but erases the advantage in SSQ reliability for classifiers relative to distances (exception: SVM).

In sum, DV-weighting ameliorated the impairment in pattern reliability due to the lossy binarization in correct and incorrect responses and is thus a recommendable procedure for classification-based RSA. Regarding the choice of the classifier for DV-weighted accuracies, we recommend WeiRD for high pattern reliability and SVM for high SSQ.

## Discussion

### Summary

We assessed and compared the reliability of dissimilarity measures for representational similarity analysis of MEG data. In brief, we found that 1) multivariate noise normalisation of the data strongly improved the accuracy of classifiers and the reliability of all dissimilarity measures, 2) distances were in general superior to classifiers in terms of pattern reliability, a difference that 3) could be largely ameliorated through decision-value weighting of classification accuracies, 4) in terms of reliability the Euclidean metric was en par with or better than the Pearson metric when correcting for condition-nonspecific response components, 5) cross-validation provided robust unbiased distance estimates for the Euclidean distance, but came at the cost of slight reliability reductions and was unstable for the Pearson distance, and 6) within-class correction addressed the problematic influence of condition-nonspecific response components on Pearson distances.

### Multivariate noise normalisation improves classification accuracies and reliability

Multivariate noise normalisation substantially improved classification accuracies and the reliability across classifiers and distance measures. Importantly, the success of noise normalisation depended on a number of important methodological details.

First, the efficacy of noise normalisation critically depended on the specific method used to compute the cross-sensor covariance matrix Σ. For instance, computing Σ just on the baseline data (*baseline method*) degraded RDM reliabilities, which may either suggest that using the baseline phase provided insufficient data for a robust estimate of Σ, or that the baseline phase was not representative of the noise characteristics during stimulus presentation. We believe that our data speak to the latter interpretation, as another method that estimated Σ separately for each time point (*time point method*) led to equal improvements in reliability. As a bottom line, we thus recommend to always include data from the stimulus period in the estimation of Σ, although great care is required to ensure that Σ is not compromised by signal information. Here, this was achieved by computing Σ for each condition separately and subsequent averaging across conditions.

Second, it should be noted that for our method of choice (*epoch method*), univariate noise normalisation was already responsible for a large gain in reliability. Thus, the increase in reliability was largely based on a univariate down-weighting of noisy sensors and up-weighting of sensors with little noise variance.

Yet third, there was nevertheless a significant advantage of MNN relative to UNN. Thus, emphasizing spatial frequencies of the MEG patterns with lower variance and de-emphasizing frequencies with higher variance (i.e., the *multiv*ariate aspect of MNN) yielded a further gain in reliability.

Overall, multivariate noise normalisation is a highly recommended preprocessing step irrespective of other analytic choices. However, the specific implementation of noise normalisation has to be chosen carefully and should consider potentially different noise structures between baseline and stimulus-driven activity.

### Choosing a classifier for decoding

When using multivariate decoding to characterize brain representations it is in most cases desirable to maximize classification accuracy. The main reason is that most neuroimaging measurements including MEG suffer from decreased sensitivity, i.e. only a fraction of the information that is encoded by the underlying neuronal ensembles can be decoded at the level of MEG or at the level of other non-invasive techniques. It is thus desirable to fully exploit every bit of information that is contained in these measurements, thereby also increasing the chance to meaningfully test more subtle comparisons between experimental conditions.

Here, we compared four classifiers (LDA, SVM, WeiRD, GNB) with and without multivariate noise normalisation and found two main results. First, whether or not to perform MNN was more critical than the choice of a classifier itself, yielding improvements in (peak) classification accuracy between 5 and 20%. Second, comparing classifiers with prior MNN, we found that LDA, SVM and WeiRD achieved all high and comparable levels of accuracy (>90% peak accuracy in our dataset), while GNB performed much worse. Overall we thus strongly advice the use of multivariate noise normalisation and recommend LDA, SVM and WeiRD as suitable classifiers for MEG decoding.

### Decision-value weighting boosts the pattern reliability of classification accuracies

Confirming previous work, we found that the pattern RDM reliability of classification accuracies was markedly impaired compared to distance measures (Walther et al., 2016). One likely reason for this clear handicap of classification accuracies is the loss of precision through the binarization of predictions (Walther et al., 2016). Another reason could be the fact that classification accuracies naturally decrease with noise and are thus not unbiased. Classification accuracies are thus not recommended for RSA.

To address these problems of classification accuracy, in the present work we introduced DV-weighting of classification accuracies as a potential solution. By weighting the correctness of single predictions with classifier decision values, gradual information is reintroduced into classification-based dissimilarity measures. Our results showed that DV-weighting indeed rectified the loss in pattern reliability for classification, advancing it close to (LDA, SVM) or to (WeiRD) the level of distance measures.

In sum, our results discourage the use of raw classification accuracies for RSA and instead advocate DV-weighting of accuracies if classification-based RSA is desired.

### Deciding between the Pearson and the Euclidean distance metric

If reliability were the only criterion, our results would strongly favour the non-cross-validated Pearson distance over the non-cross-validated Euclidean distance. However, this conclusion should be critically vetted in view of an important caveat: while the Euclidean distance is invariant to condition-nonspecific response components, the Pearson distance is strongly affected by such signals. Indeed, in most cases the Pearson distance will be a mixture of the dissimilarity due to both condition-specific signals (i.e., the signal of interest) and condition-nonspecific signals. This hampers interpretation not only of Pearson distance per se, but also of its reliability which can be inflated by condition-nonspecific response components.

Empirically, an inflation of reliability through condition-nonspecific response components was supported by two facts. First, the reliability of the Pearson metric was dramatically reduced when the mean pattern was removed prior to distance computation (cocktail-blank removal). Acknowledging that this procedure introduced new dependencies – a negative correlation – between conditions, the result conclusively indicates that the mean pattern was a driving factor for the high reliability of the Pearson distance. Second, the reliability of the Pearson distance was likewise reduced when distances were subjected to within-class correction, while this was not the case for the Euclidean distance. In both cases, the reliability of the Pearson distance became indistinguishable from the Euclidean distance to the point of being reversed in favour of the Euclidean distance. These two analyses thus strongly suggest that the high reliability of the non-cross-validated Pearson distance was partly based on condition-nonspecific response components.

Overall, in the absence of a priori reasons to prefer a mean/scale-invariant dissimilarity measure, we thus recommend the Euclidean distance over the Pearson distance for better interpretability and higher condition-specific reliability. If mean/scale invariance is desired, it is advisable to carefully check the impact of condition-nonspecific response components on the goal of RSA. In a best-case scenario, these components cancel out in a comparison of RDMs across modalities and have little effect on accuracy. However, if the contribution of condition-nonspecific components substantially differs between modalities in a condition-pair-specific manner, the accuracy of RSA might be severely impaired.

### The case for cross-validation

The unique advantage of cross-validation is that it provides unbiased estimates for distances between conditions. Indeed, unbiased distance estimates are critical for a number of research questions. Consider that we construct an RDM using the Euclidean distances measure and are interested in whether certain parts of the RDM are different from zero, i.e. we want to test whether the conditions in these parts are meaningfully different in terms of their neural representations. This test would be impossible for the non-cross-validated Euclidean distance, which is inflated by noise and produces non-zero distance estimates even for identical neural representations. Cross-validation enables testing this hypothesis by cancelling out noise between partitions of the data, thereby introducing a meaningful zero point. In a similar vein, cross-validation allows testing for ratios between distances e.g. whether distance A is twice as big as distance (Walther et al., 2016).

Despite this key advantage, our data and simulations suggest that cross-validation may come at a cost in terms of measurement reliability. While the reliability of the Euclidean distance was unchanged by cross-validation in our data, the reliability of the Pearson distance showed a significant decrease. These results imply that a potential positive effects of cross-validation on reliability (e.g., increased robustness with respect to varying noise levels between measurements), if at all present, could not outweigh the disadvantageous effects of data splitting in cross-validation. Although a negative effect of data splitting should be mitigated through averaging across cross-validation folds, this may often not be fully compensatory.

Despite this potential hit on reliability, for two reasons our results nevertheless lend support to cross-validation specifically for the Euclidean distance. First, the non-cross-validated Euclidean distance, much more than the Pearson distance, was severely distorted in the presence of noise, which in some cases rendered distance estimates almost uninterpretable. Thus, the need of cross-validation was especially pressing for the Euclidean metric. Second, cross-validation was robust for the Euclidean distance as it did not suffer from instability issues affecting the cross-validated Pearson distance (see below). Overall, cross-validation is thus a recommended procedure for the Euclidean distance.

In contrast, we recommend to forgo cross-validation for the Pearson distance for two reasons. First, the non-cross-validated Pearson distance is invariant to the level of noise in phases where the true signal is zero (e.g., baseline). This is because the expected correlation coefficient is zero in the absence of any signal, irrespective of the noise level. And second, cross-validation of the Pearson distance was associated with a consistent and often marked loss in reliability both in our data in simulation. A likely reason for this relatively strong negative impact of cross-validation on reliability are the unstable cross-validated variances which form the denominator of the cross-validated Pearson distance. These variances can easily reach close-to-zero or negative values in realistic datasets and thus dramatically distort the resulting Pearson distances. To address this instability issue, careful regularization is required. Yet, even if regularization is successful, the techniques and parameters of regularization may differ between modalities, introducing arbitrary choices and complicating cross-modal comparisons. Together, these results and considerations advise against cross-validating the Pearson distance.

### The case for within-class correction

In our data, within-class correction was slightly more reliable than cross-validation for the Euclidean distance and markedly more reliable for the Pearson distance (with cocktail-blank removal). Simulation corroborated this empirical result. Nevertheless, there are additional considerations that should guide the decision of whether to use a within-class-corrected processing scheme.

First, within-class correction provides unbiased distance estimates only for the Euclidean distance metric and only if noise affects within- and between-condition distances to the same degree. Given highly similar shapes of cross-validated (Figure 3A) and within-class-corrected (Figure 4A) Euclidean distances, this condition seemed to be met in our dataset, but might not generally be the case. Given that the reliability gains of within-class correlation relative to cross-validation are very small in the case of the Euclidean distance (ΔR_pattern/SSQ_^~^0.01), cross-validation therefore remains the recommended procedure for the Euclidean distance.

Second, in most cases within-class correction will be computationally costlier than cross-validation. The within-class-corrected algorithm applied here assessed the full permutation scheme by computing distances for all combinations of pseudo-trials within a condition and between conditions. This was feasible due to the relatively small number of pseudo-trials per condition. However, as the number of permutations grows quadratically with the number of (pseudo-)trials, random subsampling schemes will become necessary at some point, which adds algorithmic complexity and potentially a certain degree of randomness.

And third, while these first two considerations might swing the pendulum to cross-validation, there nevertheless remains the fact that only within-class correction removes condition-nonspecific signal components. This aspect is important for the Pearson distance, which, as noted above, can be strongly affected by such signals. Thus, if one wishes to use the Pearson distance without the influence of condition-nonspecific signal components and without the bias of cocktail-blank removal, within-class correction is the recommended procedure.

In sum, within-class-corrected distances come both with advantages (eliminated condition-nonspecific signal contributions, meaningful zero point, often higher reliability than cross-validation) and disadvantages (distances are not generally unbiased, increased computational complexity). We recommend to use within-class correction in cases were condition-nonspecific signal contributions are a severe issue, as in the case of the Pearson metric.

### Generalization to EEG

As the present work was focused on MVPA for MEG, to what degree can these results be generalized to MVPA for electroencephalography (EEG) data? Despite a number of physical differences between the brain’s generated magnetic and electric fields (Hämäläinen et al., 1993), a recent study by Cichy and Pantazis (2016) found largely convergent results in a comprehensive investigation of MVPA methods applied to MEG and EEG. We are therefore optimistic that the results of the present work will show good generalization to EEG data, although a specialized treatment of EEG MVPA will have to confirm this prediction.

### Conclusion and final recommendations

For both decoding and RSA we strongly recommend to apply multivariate noise normalisation on the data, which provided a considerable boost in terms of classification accuracy and reliability. By contrast, we do not recommend to use cocktail-blank removal as a preprocessing step, which failed at its primary task – removing dependencies between conditions – by introducing other dependencies, confirming earlier reports (Diedrichsen et al., 2011; Garrido et al., 2013; Walther et al., 2016).

If the goal of MVPA is decoding of MEG signals in an information-based framework (i.e., classification accuracy), we recommend to use LDA, SVM or WeiRD as classifiers, which yielded comparable accuracy. For MEG RSA, weighing in empirical results and considerations of practicability, and assuming there are no a priori constraints on the choice of a dissimilarity metric, we recommend the cross-validated Euclidean distance as a default choice, which provides a gradual, unbiased and yet reliable dissimilarity measure. Moreover, in comparison to the Pearson distance, the Euclidean distance is much less affected by condition-nonspecific signals common to different conditions and does not suffer from instability in cross-validated processing schemes. Nevertheless, when condition-nonspecific signals are not problematic or when the use of within-class correction is feasible, the Pearson distance is likewise a reliable and recommended choice for MEG RSA. If classification-based RSA is desired, we recommend to try out DV-weighting of correct and incorrect predictions, which led to considerable boosts for the pattern reliability of RDMs.

## Acknowledgements

We thank Martin N. Hebart for providing insightful comments to a previous version of the manuscript. This research was funded by an Emmy Noether Award (CI 241/1-1) of the German Research Foundation (Deutsche Forschungsgemeinschaft; DFG) to RMC, and by the DFG research group 1617, grants STE 1430/6-1 and STE 1430/6-2.

## Footnotes

1 Note that the SSQ reliability cannot be meaningfully computed for the non-cross-validated Euclidean distance and thus is not considered in this comparison.

## References

Abraham, A., Pedregosa, F., Eickenberg, M., Gervais, P., Mueller, A., Kossaifi, J., Gramfort, A., Thirion, B., Varoquaux, G., 2014. Machine learning for neuroimaging with scikit-learn. Front. Neuroinform. 8, 14. doi:10.3389/fninf.2014.00014

Chang, C., Lin, C., 2011. LIBSVM: A Library for Support Vector Machines. ACM Trans. Intell. Syst. Technol. 2, 27:1–27:27.

Cichy, R.M., Khosla, A., Pantazis, D., Oliva, A., 2017a. Dynamics of scene representations in the human brain revealed by magnetoencephalography and deep neural networks. Neuroimage 153, 346–358. doi:10.1016/j.neuroimage.2016.03.063

Cichy, R.M., Khosla, A., Pantazis, D., Torralba, A., Oliva, A., 2016a. Comparison of deep neural networks to spatio-temporal cortical dynamics of human visual object recognition reveals hierarchical correspondence. Sci. Rep. 6, 27755. doi:10.1038/srep27755

Cichy, R.M., Kriegeskorte, N., Jozwik, K.M., Van den Bosch, J.J.F., Charest, I., 2017b. Neural dynamics of real-world object vision that guide behaviour. Bioarxiv. doi:https://doi.org/10.1101/147298

Cichy, R.M., Pantazis, D., 2016. Multivariate pattern analysis of MEG and EEG: a comparison of representational structure in time and space. Bioarxiv 1–30. doi:10.1101/095620

Cichy, R.M., Pantazis, D., Oliva, A., 2016b. Similarity-Based Fusion of MEG and fMRI Reveals Spatio-Temporal Dynamics in Human Cortex During Visual Object Recognition. Cereb. Cortex 26, 3563–3579. doi:10.1093/cercor/bhwl35

Cichy, R.M., Pantazis, D., Oliva, A., 2014. Resolving human object recognition in space and time. Nat. Neurosci. 17, 455–62. doi:10.1038/nn.3635

Cichy, R.M., Sterzer, P., Heinzle, J., Elliott, L.T., Ramirez, F., Haynes, J.-D., 2013. Probing principles of large-scale object representation: category preference and location encoding. Hum. Brain Mapp. 34, 1636–1651. doi:10.1002/hbm.22020

Cox, D.D., Savoy, R.L., 2003. Functional magnetic resonance imaging (fMRI) “brain reading”: detecting and classifying distributed patterns of fMRI activity in human visual cortex. Neuroimage 19, 261–270. doi:10.1016/S1053-8119(03)00049-l

Diedrichsen, J., Ridgway, G.R., Friston, K.J., Wiestler, T., 2011. Comparing the similarity and spatial structure of neural representations: A pattern-component model. Neuroimage 55, 1665–1678. doi:10.1016/j.neuroimage.2011.01.044

Furl, N., Lohse, M., Pizzorni-Ferrarese, F., 2017. Low-frequency oscillations employ a general coding of the spatio-temporal similarity of dynamic faces. Neuroimage. doi:10.1016/j.neuroimage.2017.06.023

Garrido, L., Vaziri-Pashkam, M., Nakayama, K., Wilmer, J., 2013. The consequences of subtracting the mean pattern in fMRI multivariate correlation analyses. Front. Neurosci. 7, 1–4. doi:10.3389/fnins.2013.00174

Golarai, G., Ghahremani, D.G., Whitfield-Gabrieli, S., Reiss, A., Eberhardt, J.L., Gabrieli, J.D.E., Grill-Spector, K., 2007. Differential development of high-level visual cortex correlates with category-specific recognition memory. Nat. Neurosci. 10, 512. doi:10.1038/nnl865

Guggenmos, M., Schmack, K., Sterzer, P., 2016. WeiRD - a fast and performant multivoxel pattern classifier. 2016 Int. Work. Pattern Recognit. Neuroimaging 1–4. doi:10.1109/PRN 1.2016.7552349

Hämäläinen, M.S., Hari, R., Ilmoniemi, R.J., Knuutila, J., Lounasmaa, O. V, 1993. Magnetoencephalography - theory, instrumentation, and applications to noninvasivee studies of the working human brain. Rev. Mod. Phys. doi:10.1103/RevModPhys.65.413

Haxby, J. V, Gobbini, M.I., Furey, M.L., Ishai, A., Schouten, J.L., Pietrini, P., 2001. Distributed and overlapping representations of faces and objects in ventral temporal cortex. Science (80-.). 293, 2425–30. doi:10.1126/science,1063736

Haynes, J.-D., Rees, G., 2005. Predicting the orientation of invisible stimuli from activity in human primary visual cortex. Nat. Neurosci. 8, 686–91. doi:10.1038/nnl445

Kamitani, Y., Tong, F., 2005. Decoding the visual and subjective contents of the human brain. Nat. Neurosci. 8, 679–85. doi:10.1038/nnl444

Khaligh-Razavi, S.M., Kriegeskorte, N., 2014. Deep Supervised, but Not Unsupervised, Models May Explain IT Cortical Representation. PLoS Comput. Biol. 10. doi:10.1371/journal.pcbi.1003915

Kiani, R., Esteky, H., Mirpour, K., Tanaka, K., 2007. Object Category Structure in Response Patterns of Neuronal Population in Monkey Inferior Temporal Cortex. J. Neurophysiol. 97, 4296–4309. doi:10.1152/jn.00024.2007

Kietzmann, T.C., Gert, A.L., Tong, F., König, P., 2017. Representational Dynamics of Facial Viewpoint Encoding. J. Cogn. Neurosci. 29, 637–651. doi:10.1162/jocn_a_01070

Kriegeskorte, N., Mur, M., Ruff, D. a, Kiani, R., Bodurka, J., Esteky, H., Tanaka, K., Bandettini, P. a, 2008. Matching categorical object representations in inferior temporal cortex of man and monkey. Neuron 60, 1126–41. doi:10.1016/j.neuron.2008.10.043

Ledoit, O., Wolf, M., 2004. Honey, I Shrunk the Sample Covariance Matrix. J. Portf. Manag. 30, 110–119. doi:10.3905/jpm.2004.110

Mur, M., Meys, M., Bodurka, J., Goebel, R., Bandettini, P.A., Kriegeskorte, N., 2013. Human object-similarity judgments reflect and transcend the primate-IT object representation. Front. Psychol. 4, 1–22. doi:10.3389/fpsyg.2013.00128

Myers, N.E., Rohenkohl, G., Wyart, V., Woolrich, M.W., Nobre, A.C., Stokes, M.G., Prlic, A., Wacongne, C., Labyt, E., Bekinschtein, T., Cohen, L., Naccache, L., Dehaene, S., 2015. Testing sensory evidence against mnemonic templates. Elife 4, e09000. doi:10.7554/eLife.09000

Op de Beeck, H.P., 2010. Against hyperacuity in brain reading: Spatial smoothing does not hurt multivariate fMRI analyses? Neuroimage. doi:10.1016/j.neuroimage.2009.02.047

Pantazis, D., Fang, M., Qin, S., Mohsenzadeh, Y., Li, Q., Cichy, R.M., 2017. Decoding the orientation of contrast edges from MEG evoked and induced responses. BioRxiv. doi:https://doi.org/10.1101/148056

Pietrini, P., Furey, M.L., Ricciardi, E., Gobbini, M.I., Wu, W.-H.C., Cohen, L., Guazzelli, M., Haxby, J. V, 2004. Beyond sensory images: Object-based representation in the human ventral pathway. Proc. Natl. Acad. Sci. U. S. A. 101, 5658–63. doi:10.1073/pnas.0400707101

Seeliger, K., Fritsche, M., Güçlü, U., Schoenmakers, S., Schoffelen, J.-M., Bosch, S.E., Van Gerven, M.A.J., 2017. CNN-based Encoding and Decoding of Visual Object Recognition in Space and Time. BioRxiv. doi:https://doi.org/10.1101/118091

Su, L., Fonteneau, E., Marslen-Wilson, W., Kriegeskorte, N., 2012. Spatiotemporal searchlight representational similarity analysis in EMEG source space. Proc. - 2012 2nd Int. Work. Pattern Recognit. NeuroImaging, PRNI 2012 97–100. doi:10.1109/PRNI.2012.26

Walther, A., Nili, H., Ejaz, N., Alink, A., Kriegeskorte, N., Diedrichsen, J., 2016. Reliability of dissimilarity measures for multi-voxel pattern analysis. Neuroimage 137, 188–200. doi: 10.1016/j.neuroimage.2015.12.012

Wardle, S.G., Kriegeskorte, N., Grootswagers, T., Khaligh-Razavi, S.M., Carlson, T.A., 2016. Perceptual similarity of visual patterns predicts dynamic neural activation patterns measured with MEG. Neuroimage 132, 59–70. doi:10.1016/j.neuroimage.2016.02.019

Weiner, K.S., Sayres, R., Vinberg, J., Grill-Spector, K., 2010. fMRI-adaptation and category selectivity in human ventral temporal cortex: regional differences across time scales. J. Neurophysiol. 103, 3349–65. doi:10.1152/jn.01108.2009

Williams, M.A., Baker, C.I., Op de Beeck, H.P., Mok Shim, W., Dang, S., Triantafyllou, C., Kanwisher, N., 2008. Feedback of visual object information to foveal retinotopic cortex. Nat. Neurosci. 11, 1439–1445. doi:10.1038/nn.2218

Williams, M.A., Dang, S., Kanwisher, N.G., 2007. Only some spatial patterns of fMRI response are read out in task performance. Nat. Neurosci. 10, 685–686. doi:10.1038/nn1900

